# Protection of algae grown for biofuel using a consortium of environmentally harvested bacteria

**DOI:** 10.64898/2026.03.18.712687

**Authors:** Elise K. Wilbourn, Deanna Curtis, John McGowen, Pamela Lane, Everett Eustance, Olivia Watt, Tyler P. Eckles, Todd W. Lane

## Abstract

Crop loss due to infection by pests and pathogens is a major barrier to the large-scale production of algal biofuels. Test systems have seen loss of green algae crops due to infection by the fungus-like *Amoeboaphelidium occidentale* FD01. While current antifungal compounds are effective in inhibiting the infection, their application raises the overall cost of the crop and lowers its economic viability as a biofuel source. Here we show that co-culturing environmentally harvested bacteria alongside algae crops can drastically lower the rate of infection in two different green algae species of interest for biofuel production. These bacteria-algae consortia increase the mean time to crop failure (MTTF) by up to 350% when tested under environmentally relevant conditions. While there was an increase in diversity over time, there was no statistically significant correlation between an increase in diversity and a longer MTTF. Community composition analysis reveals similarities between the bacterial genera growing alongside both green algae species even as bacterial harvest locations differed, although there was not a single dominant genus responsible for the increase in crop protection. These results show a promising new method of anti-fungal crop protection that can be applied to algal biofuels with no increase in fuel cost.

**Highlights:**

- Bacteria-algal cocultures protect against fungal pests without impact to productivity
- Bacterial community composition is variable over time even as protection persists
- Bacterial consortia can increase mean time to failure by 350%

## 1 Introduction

Algae are a promising source of many commercial products, including biofuels, proteinaceous biomass, and high-value nutraceutical products [1–4]. To reduce production costs, planktonic algae can be grown in large-scale open-air outdoor ponds at volumes of 100,000 liters or higher [5]. As the ponds are generally open to the elements, unialgal crops are susceptible to biotic contaminants, including infection by pests [6, 7]. Infection can lead to a reduction in crop productivity that has been estimated as to be as high as 30% [8, 9]. As the base of the aquatic food system, algae are susceptible to predation by a variety of organisms. Pests present in algae ponds can include bacteria, viruses, grazing zooplankton, and fungus. One group of interest due to their apparent prevalence in outdoor algae ponds is the aphelids. The phylum *Aphelidea* is a poorly understood member of the group *Opisthokonta* related to fungi [10–12]. An infection by a single species of aphelid can cause complete population crash in a pond of green algae in only a few days. One of the earliest described aphelid species found in algal ponds is *Amoeboaphelidium protococcarum* [13, 14]. All phyla of aphelids including amoeboaphelids tend to be highly host-specific, infecting only one or a few species and almost always infecting green algae (although there is at least one known species capable of infecting a diatom species [15]). Letcher et al. [14] saw infection of a green algae species, *Scenedesmus dimorphous*, commonly cultured for biofuels, by *Amoeboaphelidium protococcarum* strain FD01. Later studies confirmed the ability of FD01 to infect other *Scenedesmus* species [16], with the potential to infect other green algae species[17].

Current methods for dealing with fungal contaminants include immediate harvesting of the algal crop as soon as the infection is detected and/or application of either prophylactic or post-infection antifungal compounds. However, both methods nearly always result in a loss of crop and lower product yield, increasing the cost of the end products. Especially in the case of biofuels and other low-value per volume products, this can raise the cost of algae-based products far above the cost of conventional fossil-fuel based products [7, 18]. Additionally, widespread application of antifungals or other broad-spectrum antibiotics risks the development of antibiotic-resistant fungal species as has been seen in human and plant fungal pests [19–21]. There are a limited number of antifungal compounds currently approved for use in the United States, with several dozen compounds restricted at the federal level and more regulations on a state-by-state basis, while in the European Union each member country must individually approve plant protection products including antifungals. Currently used antifungal compounds used in algal biofuel crops include fluconazole and other azole compounds [22], rotenone [23–25], fluazinam, and quinine [26]. These compounds often show anti-protist activity in addition to their anti-fungal properties, lowering the final algal biomass and product yield and therefore further raising the cost of the end products.

In natural systems algae often grow alongside highly diverse bacterial assemblages, and these bacteria interact with the algae in a variety of ways. This interaction can range from beneficial to the algae to deleterious [27–29], but generally the interaction between the two populations shows no negative effect on any member. The scale needed to product economically viable algal biofuels is such that a closed system is nearly impossible, so it should be expected that similar bacterial populations will be present in algal production ponds. In terrestrial crop systems, modification of the microbiome with beneficial bacterial species has been shown to increase total crop yield [30–32]. This modification can range in price, but often the final increase in yield often makes it an economically viable method. In addition to potentially increasing crop yield, modification of the microbiome in algal systems has been attempted as a method of combatting crop loss [33]. In terrestrial crops, application of beneficial bacteria must be repeated on a semi-regular basis due to the inherent instability of the soil microbiome, which responds to environmental changes across the course of the growing season.

Less work has been done regarding modification of biofuel algae crop microbiomes. Our study focuses on two algal species: *Monoraphidium minutum* strain 26B-AM and *Tetradesmus obliquus* strain UTEX393 (previously *Scenedesmus obliquus*). These species have been previously studied as possible sources of algal biofuel [34] due to their fast growth rate and high lipid production levels. They have been grown successfully in both laboratory-based closed systems and in outdoor pilot- and production-scale systems, and the optimal growth conditions are known. Past studies have shown that the associated bacteria are capable of providing anti-rotifer protection [35] but no work has been done on using an environmental bacteria-algae consortia for protection against fungal pests in either species. Here we characterize the microbiome of two algal cultures following inoculation with environmental bacterial consortia over multiple months. Additionally, we demonstrate the ability of two consortia to provide protection against a fungus-like pathogen, *Amoeboaphelidium occidentale* FD01 [13].

## 2 Materials and Methods

### 2.1 Culturing

The algal species were cultured in enclosed, lighted, and temperature-controlled incubators or environmental photobioreactors (ePBRs) for all experiments. Cultures in incubators were grown in 14:10 hour light:dark cycle with 150 µE illumination. Algae were cultured in either 50 mL sterile polypropylene tubes (VWR) containing 20-25 mL of culture or 500 mL sterile glass baffled Erlenmeyer flasks containing 120-125 mL culture and closed with vented aluminum caps. Both tubes and flasks were continuously shaken at approximately 100 rpm. Cultures were diluted approximately every three weeks by diluting 1:25 vol/vol with sterile algal culturing medium to prevent nutrient stress.

### 2.2 Algal Culturing Medium

Artificial seawater salts were prepared according to [36, 37] and diluted with DI water to 5 parts per thousand (ppt) total salts. Nutrients were then added according to a previously optimized recipe [38] at the following concentrations: (NH_4_)_2_SO_4_ at 4.5 mM, (NH_4_)_2_HPO_4_ at 0.27 mM, CaCl_2_.2H_2_O at 0.024 mM, MgSO_4_.7H_2_O at 0.032 mM, NaHCO_3_ at 3.572 mM, and KCl at 0.134 mM. Micronutrients and trace metals were added following the recipe for f/2 trace metals and micronutrient solutions [39, 40]. Once all salts and nutrients were dissolved the media was filtered into a sterile bottle through a sterile filter tower (0.2 µm pore size, PTFE, VWR).

### 2.3 Environmental Photobioreactors

To test algal and bacterial consortia growth in under simulated environmental conditions, cultures were grown at 500 mL scale in ePBRs (Algae Metrics)[41]. Culturing vessels were autoclaved prior to filling with 400 mL sterile algal media and 100 mL algal culture. Culture health was monitored with pH and temperature probes. Cultures were lit at 1500 µE for 14 hours light: 10 hours dark and temperature ranged sinusoidally between 32 °C at the time of peak illumination to 24 °C during the peak of the dark cycle. Cultures were both stirred with a magnetic stir bar at the bottom of the culture vessel at 120 rpm and sparged with filtered air (0.8 μm, PTFE filter, VWR). This prevented algae from settling to the bottom of the culture vessel. Samples were collected with sterile syringes through sampling stopcocks attached to each culture vessel, negating the need to remove the culture from the heating system or lights. Samples were diluted periodically to prevent nutrient stress by removing approximately 300 mL of culture and replacing this with 300 mL sterile culture (dilution times noted in figure 6).

### 2.4 Amoeboaphelid Culturing

As aphelids are obligate pests who cannot be cultured without an algal host, they were maintained through regular reinfection of unbacterialized *M. minutum* cultures. Following algal culture collapse, amoeboaphelid cultures were kept in the dark at 4 °C in sterile 50 mL polystyrene tubes (VWR) for no more than six months. Aphelids were regularly tested for infectivity using healthy, unbacterialized algal cultures.

### 2.5 Bacterial Harvesting and Inoculation

Environmental bacteria were harvested from eight locations by placing water samples into polystyrene bottles. A complete list of harvesting conditions is shown below in Table 1. Consortia A, B, J, K, L, M, and X were harvested from unialgal cultures containing the algae listed, while the others contained an undefined mixture of environmental algae.

**Table 1:** Harvesting conditions for environmental bacteria used to inoculate algal cultures (*three CA locations).

| Consortium Name | Sampling Location* | Description | Initial Algae Species |
| --- | --- | --- | --- |
| A | AZ | Outdoor algae pond | 26B-AM |
| B | AZ | Outdoor algae pond | UTEX393 |
| C | CA | Algal culture | Mixed |
| D | HI | Algal culture | Mixed |
| E | CA | Algal culture | Mixed |
| F | CA | Creek | Mixed |
| G | CA | Indoor algae pond | Mixed |
| H | CA | Indoor algae pond | Mixed |
| J | AZ | Outdoor algae pond | 26B-AM |
| K | AZ | Outdoor algae pond | 26B-AM |
| L | AZ | Outdoor algae pond | 26B-AM |
| M | AZ | Outdoor algae pond | UTEX393 |
| N | CA | Creek | Mixed |
| O | CA | Flower vase | Mixed |
| P | CA | Indoor algae pond | Mixed |
| Q | CA | Indoor algae pond | Mixed |
| R | CA | Indoor algae tub | Mixed |
| S | CA | Indoor algae pond | Mixed |
| T | CA | Algal culture | Mixed |
| U | CA | Algal culture | Mixed |
| V | CA | Indoor algae pond | Mixed |
| W | CA | Indoor algae pond | Mixed |
| X | CA | Algal culture | 26B-AM |

**Table 2:**
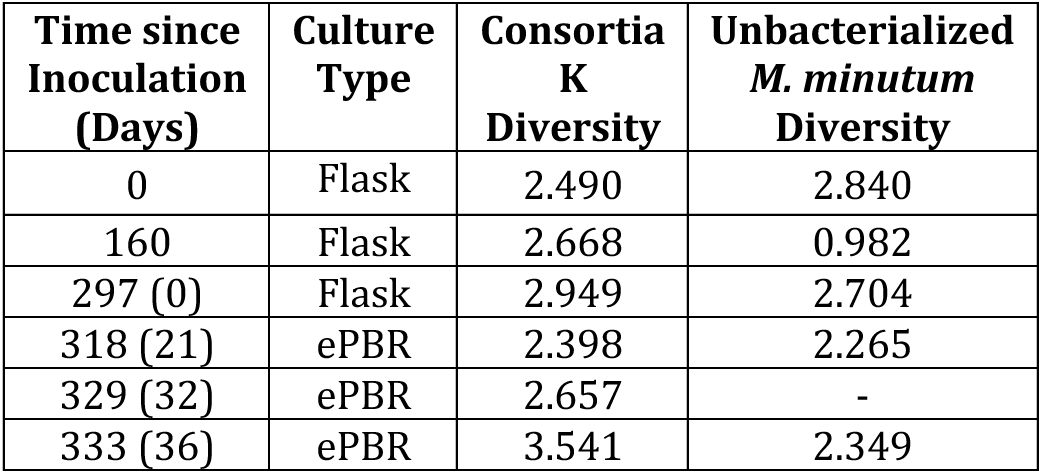
Shannon diversity over time in bacterialized and unbacterialized cultures of 26B-AM. For cultures grown in ePBRs, the time since the culture was placed into the ePBR is indicated in parentheses in the first column.

| Time since Inoculation (Days) | Culture Type | Consortia K Diversity | Unbacterialized *M. minutum* Diversity |
| --- | --- | --- | --- |
| 0 | Flask | 2.490 | 2.840 |
| 160 | Flask | 2.668 | 0.982 |
| 297 (0) | Flask | 2.949 | 2.704 |
| 318 (21) | ePBR | 2.398 | 2.265 |
| 329 (32) | ePBR | 2.657 | - |
| 333 (36) | ePBR | 3.541 | 2.349 |

**Table 3:** Shannon diversity over time in bacterialized and unbacterialized cultures of UTEX393. For cultures grown in ePBRs, the time since the culture was placed into the ePBR is indicated in parentheses in the first column.

| Time since Inoculation (Days) | Culture Type | Consortia B Diversity | Unbacterialized UTEX393 Diversity |
| --- | --- | --- | --- |
| 0 | Flask | 3.549 | 1.001 |
| 130 | Flask | 2.731 | 3.148 |
| 269 | Flask | 3.026 | 1.899 |
| 297 (0) | ePBR | 2.892 | 0.841 |
| 316 (19) | ePBR | 3.218 | 3.328 |
| 333 (36) | ePBR | 3.045 | 1.698 |

Bacteria were harvested from the water samples by filtration through a 0.8 µm pore size filter unit into a sterile tube (VWR) and the collected filtrate was kept at 4°C until it was added to algae. Each bacterial consortium was added to unialgal cultures of 26B-AM and UTEX393 and immediately tested for anti-aphelid activity. Following inoculation, bacterial and algal co-cultures were cultured according to methods described in Section 3.1. The algal cultures are referred to as bacterialized following their inoculation with bacterial consortia, but unbacterialized cultures are not axenic and contain environmental bacteria.

### 2.6 Amoeboaphelid Challenge Tests

Amoeboaphelid infectivity was tested by infected algal cultures with *A. occidentale* FD01 stock. Amoeboaphelids cannot be readily separated from crashed algal cultures, so the amoeboaphelid stock contains primarily but not exclusively amoeboaphelids. To ensure equally distributed amoeboaphelids, the crashed cultures are diluted 1:200 v/v prior to dosing create a diluted amoeboaphelid stock. Infections are tested in sterile, lidded polystyrene 48-well plates (VWR). Cultures were always tested in biological triplicate. Each well was filled with 600 µL filtered culture media (as described in Section 2.2), 300 µL algal culture, and 100 µL diluted amoeboaphelid stock, giving a final amoeboaphelid dilution of 1:2000 v/v. Plates are closed with sterile lids and stored inside a closed incubator under the same conditions described in section 4.2 but without shaking. Biomass was measured using chlorophyll *a* fluorescence as a proxy using a fluorescent plate reader (Tecan) at excitation/emission 450/685 nm. Each well was resuspended with a sterile pipette tip prior to reading fluorescence as algae settle out over time.

### 2.7 Amplicon Sequencing

Samples were stored in 15 mL sterile polypropylene tubes (VWR) in the dark at −20 °C until DNA was extracted. Samples were kept on ice where possible for the entire protocol and stored at 4°C during the library preparation process when not in use. Prior to extraction, samples were thawed and filtered through 96-well filter plates (Pall, 1 mL-0.2 µm) using centrifugation at 2500 xG to remove the brackish media from the algal and bacterial cells. The culture was resuspended from the filter using Zymo Bead Beating Buffer and DNA was subsequently extracted using the Zymo Quick-DNA Fungal/Bacterial 96 Kit according to manufacturer instructions. Extracted DNA was quantified with the QuantIt dsDNA High Sensitivity Kit (Invitrogen) and normalized in a sterile 96-well semi-skirted PCR plate (BioRad) to 2 ng/uL DNA per well. To normalize, samples were diluted with molecular-grade water (Sigma Aldrich). The V3-V4 region of the 16S gene was amplified with PCR with the plastid-excluding primers 357F and 783R [42] using 22.5 µL AccuPrime Pfx Supermix (Invitrogen), 0.5 µL each 12 µM 357F and 783R, and 6.5 µL of 2 ng/µL template DNA per well. The PCR protocol is available in the Supplemental Information and was completed on a model T100 thermal cycler (BioRad).

Following amplification samples were processed with the Zymo Clean and Concentrate 96 kit following manufacturer instructions. Samples were indexed with Nextera XT Indices (Illumina, Sets A and D). Each well contained 12.5 µL Failsafe Buffer E (BioSearch Technologies), 0.25 µL Failsafe Enzyme (BioSearch Technologies), 4.75 µL nuclease-free water (VWR), 2.5 µL of each index primer, and 2.5 µL DNA. The PCR protocol is included in the SI. Following indexing samples were cleaned according to manufacturer instructions using the same kit used previously. DNA content was then quantified using the same Quant-IT kit and the final library was prepared by combining 25 ng of DNA from each well into a single sterile polypropylene tube (VWR). The final library was quantified with the Qubit dsDNA Quantification Assay Kit (Invitrogen) and diluted to 4 nM. Past analysis of library length using a BioAnalyzer (Agilent) has shown an average library size of 609 bp, so the concentration was calculated using this assumed length. The final library was combined with 10% v/v 4 nM PhiX (Illumina) as a library control. It was then denatured to prepare for sequencing according to manufacturer instructions for the MiSeq 600 cycle Reagent Kit V3 (Illumina) and sequenced with an Illumina MiSeq.

### 2.8 Bioinformatics and Statistics

Amplicon data files from 16S sequencing were processed with Kraken 2 version 2.1.3 and reanalysis was completed with Bracken version 2.9 [43–45]. Files were compared with the Silva ribosomal RNA database version 138.1 [46, 47]. All data processing occurred in Python version 3.9.12, including calculation of the Shannon diversity index [48]. Figures were also created in Python.

## 3 Results and Discussion

When bacterial consortia were initially added to unialgal cultures and tested for protective ability against FD01, one consortium in particular (referred to as Consortium K) showed a clear antifungal ability in both algal species, although the protection was higher in 26B-AM cultures. As the bacteria and algae were incubated together, another consortium began to show antifungal activity in UTEX393, referred to as Consortium B. This consortium is not protective in 26B-AM even after 11 months of co-culturing.

### 3.1 Initial Bacterial Inoculum Compositions and Diversity

Figure 1 shows the initial bacterial community compositions inoculated into the unialgal cultures (algal culture bacterial community composition is shown in the SI). Consortium K and consortium B originated in outdoor algae pond at the Arizona State University AzCATI facility. The source pond for consortium K contained 26B-AM and the source pond for consortium B contained UTEX393. Neither pond showed signs of aphelid infection either microscopically or via community composition analysis at the time of harvest. Although the pond was filled with unialgal culture, due to the outdoor location and the exposed nature of the pond, axenic growth is not possible. Bacteria may be introduced to the pond from environmental sources including both direct interaction of the pond with insects and other local fauna and via airborne deposition. Airborne bacteria are generally transported in association with dust, pollen, or other large aerosols [49, 50], with transport distances upwards of several thousand kilometers [51, 52].

**Figure 1:**
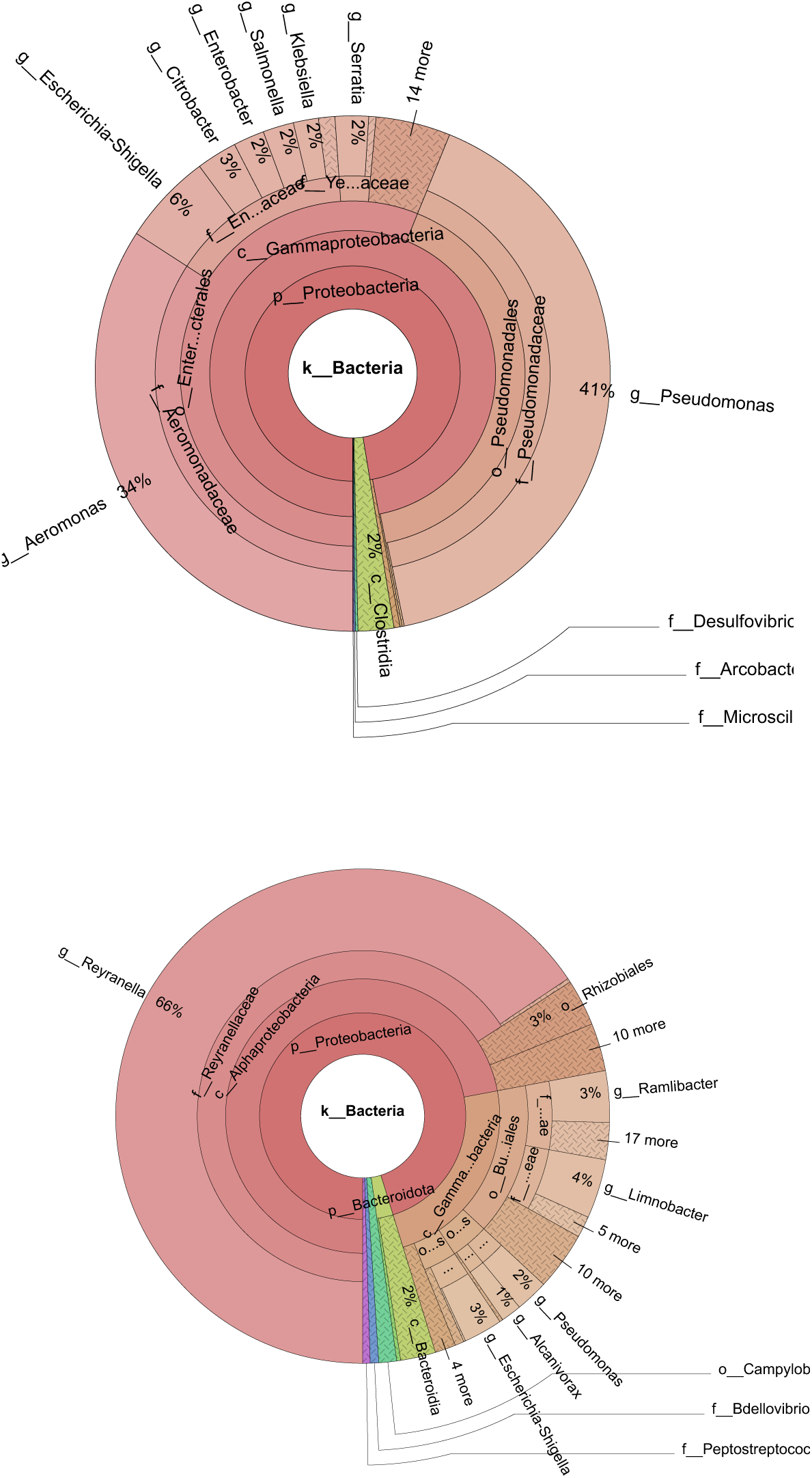
Initial bacterial community composition of protective Consortium K (above) and Consortium B (below).

The composition of the two initial consortia were similar only in their dominance by proteobacteria, with 97% of the reads in consortium K and 95% in consortium B belonging to this phylum. As the sequencing process includes two PCR amplification steps, these proportions should be considered non-quantitative but can be qualitatively compared amongst samples prepared using the same methods by comparing the relative abundance. Rather than indicating a protective consortium, a bacterial community dominated by proteobacteria may instead be a characteristic commonly seen in algae ponds or alongside green algae. When other levels of taxonomic classification are considered, there is little similarity between the two consortia. Consortium K is dominated by two genera, *Pseudomonas* (41%) and *Aeromonas* (34%) while consortium B is dominated by the genus *Reyranella* at 66% of the reads.

While all three genera have been described growing in association with algal cultures [53–55], none of the genera have been shown to have antibacterial properties when grown alongside algae. Instead, *Pseudomonas* and *Aeromonas* are commonly assumed to contain pathogenic antialgal species [56–59]. Since these consortia were not collected from ponds with lowered algal productivity, it is unlikely that the *Pseudomonas* or *Aeromonas* species contained in consortium K produce antialgal compounds. *Reyranella* is not a known antialgal or antibacterial genus. A limitation of community composition analysis via amplicon sequencing is an inability to identify organisms at the species level, which could provide more clarity on the bacterial strains responsible for protection. Additionally, initial consortia may not be dominated by the protective bacteria, so data from the bacterialized algae cultures shown below in section 5.1.1 may provide more information on the species responsible.

#### 3.1.1 Bacterialized Algae Cultures

##### 3.1.1.1 Composition Patterns in Initial Algal and Bacterial Cultures

Although some of the bacterial consortia were inoculated into the same algal strains they were harvested from, the bacterial community composition shifted somewhat after the algae were inoculated. This is expected due to a combination of the non-axenic nature of the starting algae cultures and the change in environmental conditions as the bacteria were harvested from outdoor cultures and added to flask cultures grown in enclosed incubators with lower light levels. However, the dominant class in each protective consortium also remains the same, with 62% of the class-level bacterial sequences from gammaproteobacteria in the 26B-AM culture post-inoculation (compared with 97% in the inoculum). Figures 2 and 3 show the bacterial composition of the bacterialized algae after growing with the bacterial consortia for one month.

**Figure 2:**
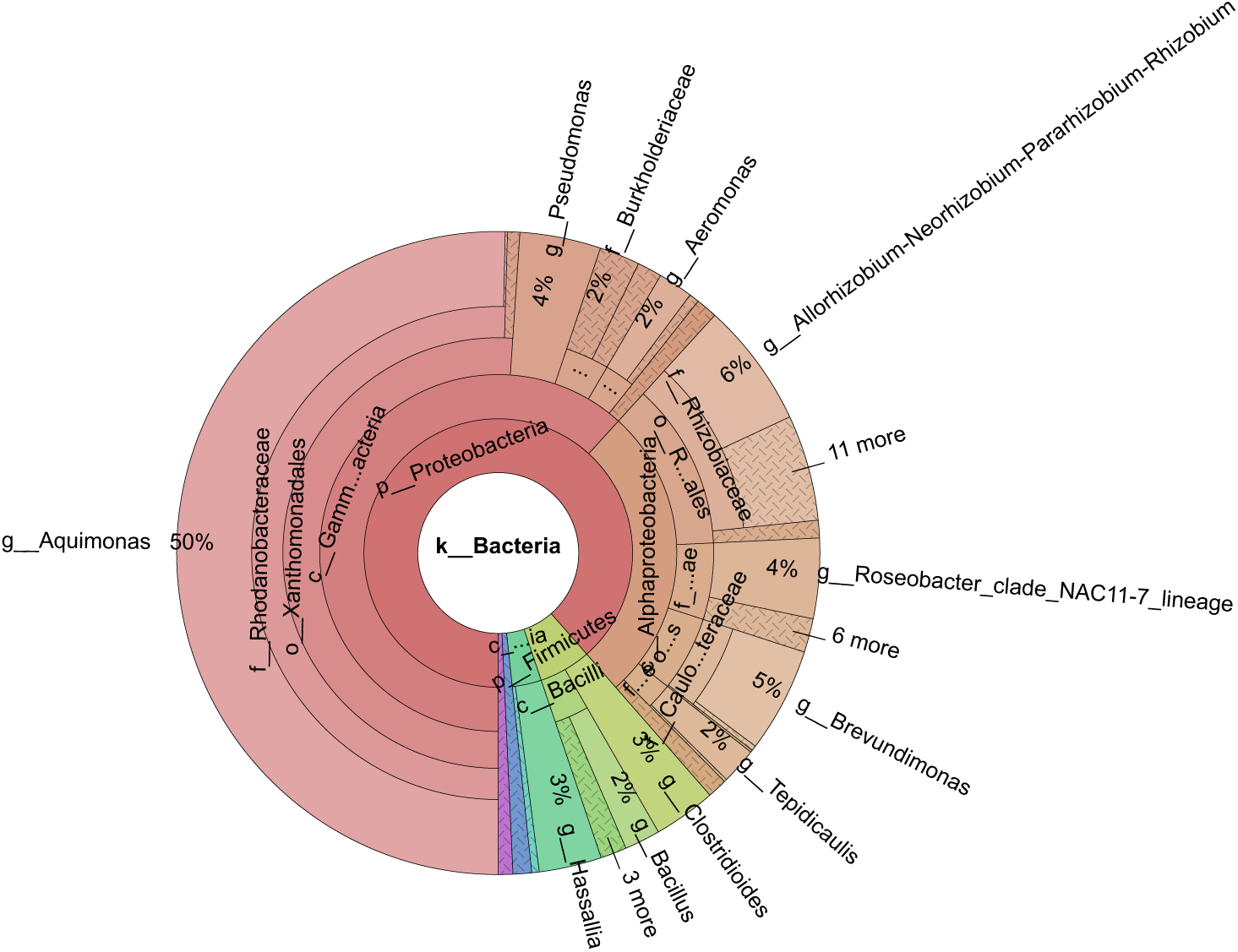
Bacterial community composition of mixed 26B-AM and bacterial consortium K after approximately one month of co-culture

**Figure 3:**
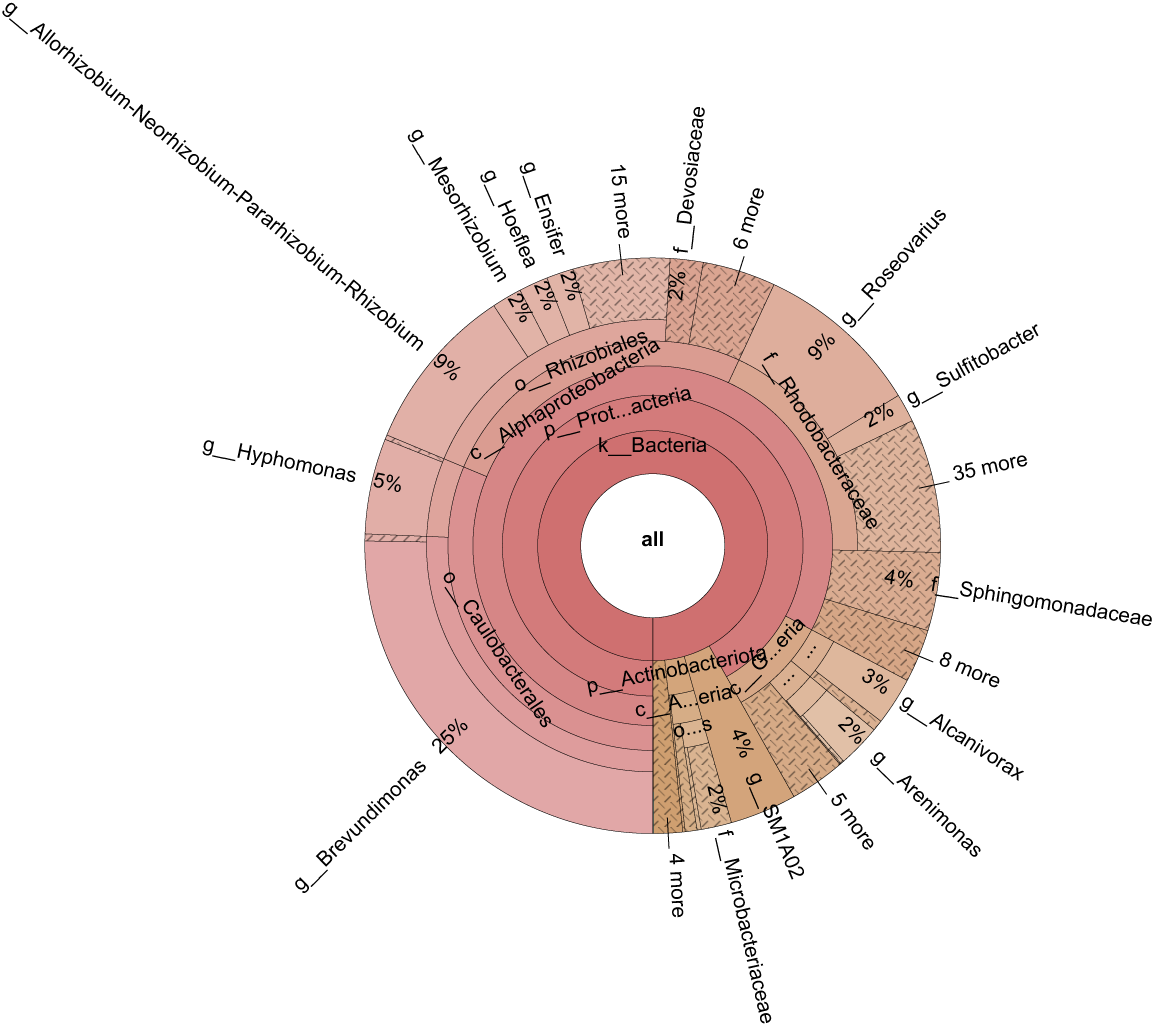
Bacterial community composition of combined UTEX393 and bacterial consortium B after approximately one month of co-culture.

Like the initial consortia, the bacteria and algae consortia are dominated by proteobacteria (89% of reads in the 26B-AM culture and 92% in the UTEX393 culture). Additionally, when the community composition of bacterialized cultures that show no change in the aphelid infection rate is compared to those of protective bacterial consortia, the non-protective consortium is also dominated by proteobacteria (see Supplemental Information for a representative example of a non-protective consortium community composition). The UTEX393 bacterialized culture contained 83% alphaproteobacteria (an increase from 72% in the inoculum). At order level and lower there is no pattern in bacterial community composition between pre- and post-inoculation cultures, although many genera are present in both samples.

### 3.2 Current Bacterial Consortia Compositions and Diversity

The bacteria and algae co-cultures were maintained for more than eight months with samples collected for sequencing. Previous studies have indicated that an increase in diversity comes alongside an increase in predator resistance [35], although this has never been shown during amoeboaphelid infection. The ending composition of the two bacterial consortia are addressed separately below.

#### 3.2.1 26B-AM

The Shannon diversity of unbacterialized and bacterialized cultures of 26B-AM did not show a large difference between most samples, indicating that unlike previous studies the protective bacterial consortium was not more diverse than the microbiome present alongside the unbacterialized culture. Although the dominant taxa changed drastically over time, as the protection was maintained, so was the diversity level. There is no pattern in diversity over time in either the bacterialized or unbacterialized samples. There is also no difference in the cultures grown in flask versus those grown in ePBRs, although this is not unexpected as the major difference between the two is the light level each culture is exposed to and the volume of culture.

The lowest diversity level in bacterialized cultures was measured in the samples grown for 21 days in the eBPR, which had a diversity of 2.398. The lowest diversity in the unbacterialized samples was much lower at 0.982 and occurred in samples taken after 160 days of culturing. This is much lower than the rest of the samples, which have generally more than twice the level of diversity.

To determine whether the bacteria continued to protect the algae from aphelid infection, 26B-AM cultures were challenged with FD01. The bacterialized algae consistently showed some protection against infection but never showed greater than 100% increase in MTTF. Once the cultures were placed into ePBRs this protection increased over time. This consistent protection in both flasks and ePBRs indicates the persistence of the protective bacteria alongside the algae.

#### 3.2.2 UTEX393

Bacterialized UTEX393 showed slightly lower resistance to infection than bacterialized 26B-AM, as well as a higher range in Shannon diversity between bacterialized and unbacterialized cultures.

This may be due to differences in the bacterial consortia between the two cocultures or due to differences in host susceptibility to infection by FD01. Unlike 26B-AM, the bacterialized UTEX393 cultures showed much higher bacterial diversity than unbacterialized cultures in more than half the measured samples in both flask-grown and ePBR samples (shown in Table 4).

### 3.3 Algal culturing in ePBRs

To determine whether the bacterial consortia were both stable during long term culturing and whether protection was maintained during growth during simulated environmental conditions, bacterialized cultures were grown in ePBRs for 30 days and tested for protection against amoeboaphelids during and following the 30 day growth period. Growth conditions are described in Section 2.3. Chlorophyll content and cell counts are shown below in Figure 4.

**Figure 4:**
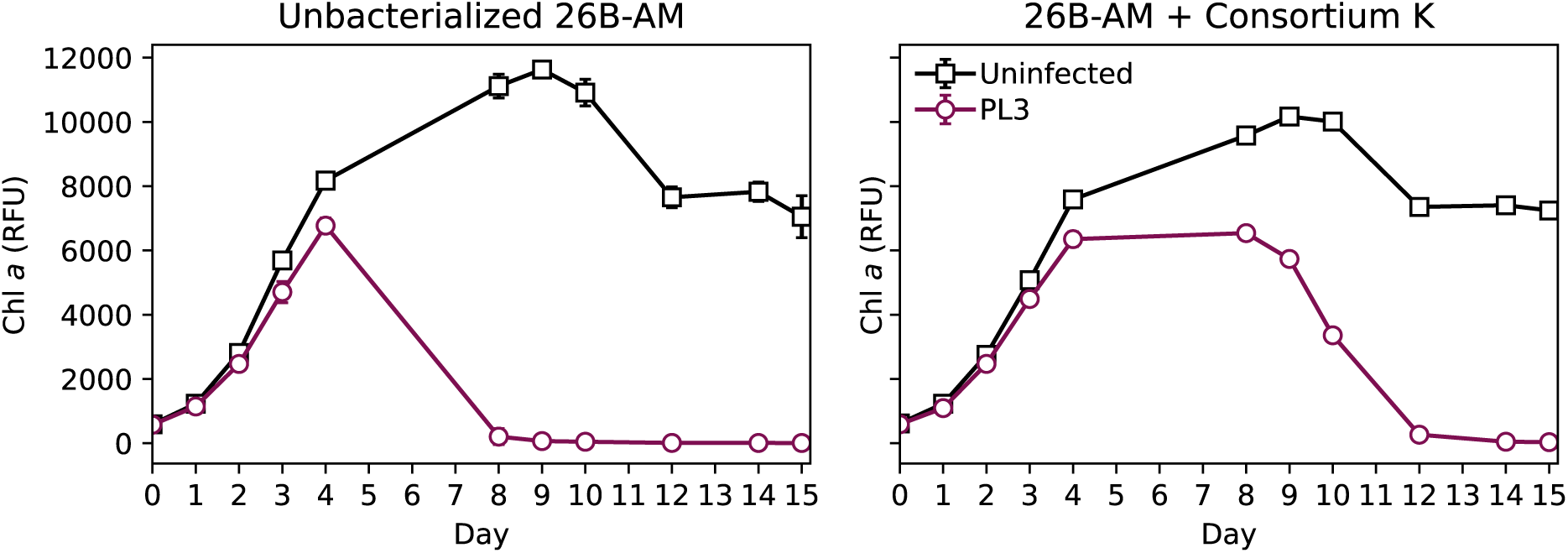
Unbacterialized and bacterialized M. minutum 26B-AM cultures infected with Amoeboaphelium sp. FD01. Data points represent the mean chlorophyll a fluorescence (RFU) +/- standard deviation (n=3).

The cultures were diluted, and fresh media was added after no more than 12 days of growth, ensuring that nutrient stress was not a factor for the algae during the infection process. Algal cell counts were diluted to approximately 10^5^ to 10^6^ mL^-1^ Although the bacterialized cultures have a higher cell concentration for the last 10 days of growth, this was not consistent during the beginning of the experiment. Additionally, the chlorophyll a fluorescence which can be used as an additional proxy for total algal biomass does not show as clear a pattern. Although there are bacterial species that can increase algal growth [60–62], it is not clear that the increase in biomass here is due to algal-bacterial interactions.

Protection by bacteria was tested by conducting amoeboaphelid challenge infections in plates using cultures removed from the ePBRs. Once the algal cultures were placed into the ePBRs, samples were taken to challenge the protection of the algae by the bacteria against the amoeboaphelids on days 11, 18, 26, and 30 of growth. The results from day 30 are shown below in Figure 5, and the results on days 11, 18, and 26 are shown in the supplemental information.

**Figure 5:**
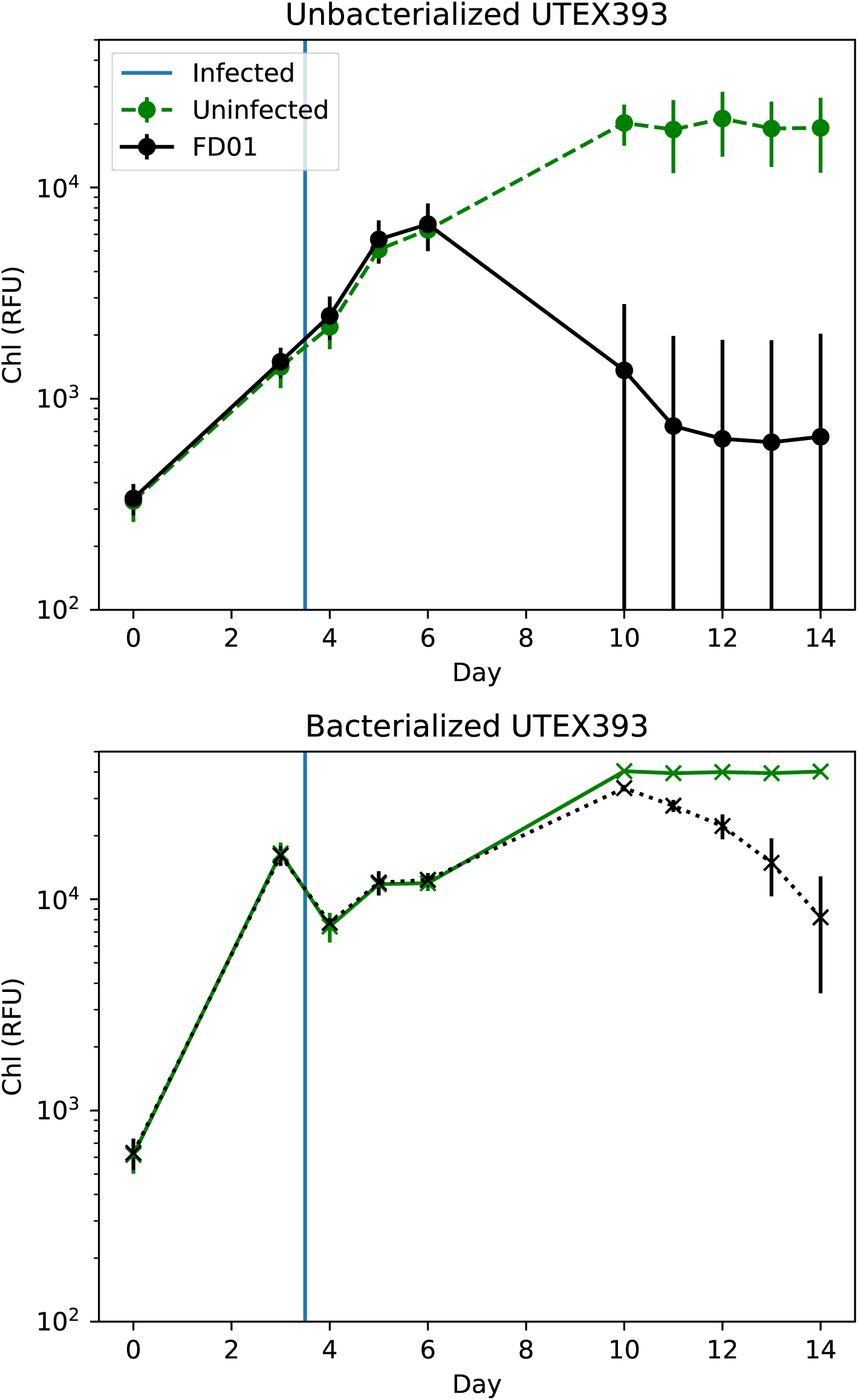
Infection of bacterialized and unbacterialized T. obliquus UTEX393 with Amoeboaphelidium sp. FD01. Data points represent the mean chlorophyll a fluorescence (RFU) +/- standard deviation (n=3).

It is clear that the protection maintains during culturing, but more interestingly as the algae and bacteria are grown together at the higher light and temperature conditions the MTTF increases in bacterialized cultures but not in unbacterialized control cultures, indicating that the protection is not due to changes in the algal cultures but is due to the bacteria. The protection is higher in the UTEX393 cultures. Although the MTTF in unbacterialized UTEX393 cultures is longer than in unbacterialized 26B-AM, the increase in MTTF by day in bacterialized UTEX393 is greater over time. However, the overall percent increase in MTTF was highest in the bacterialized 26B-AM culture, with a final increase between flask and plate MTTF of 350%.

#### 3.3.1 Community composition of protective consortia

The bacterialized cultures which showed the greatest increase in MTTF were also sequenced to look for similarities between protective consortia in both initial samples and ending samples.

Figure 6 shows the bacterial community composition of the most protective consortium grown alongside 26B-AM. When compared to both the starting bacterial consortium and the initial bacterialized culture, both the increase in diversity and the differences in composition are apparent.

**Figure 6:**
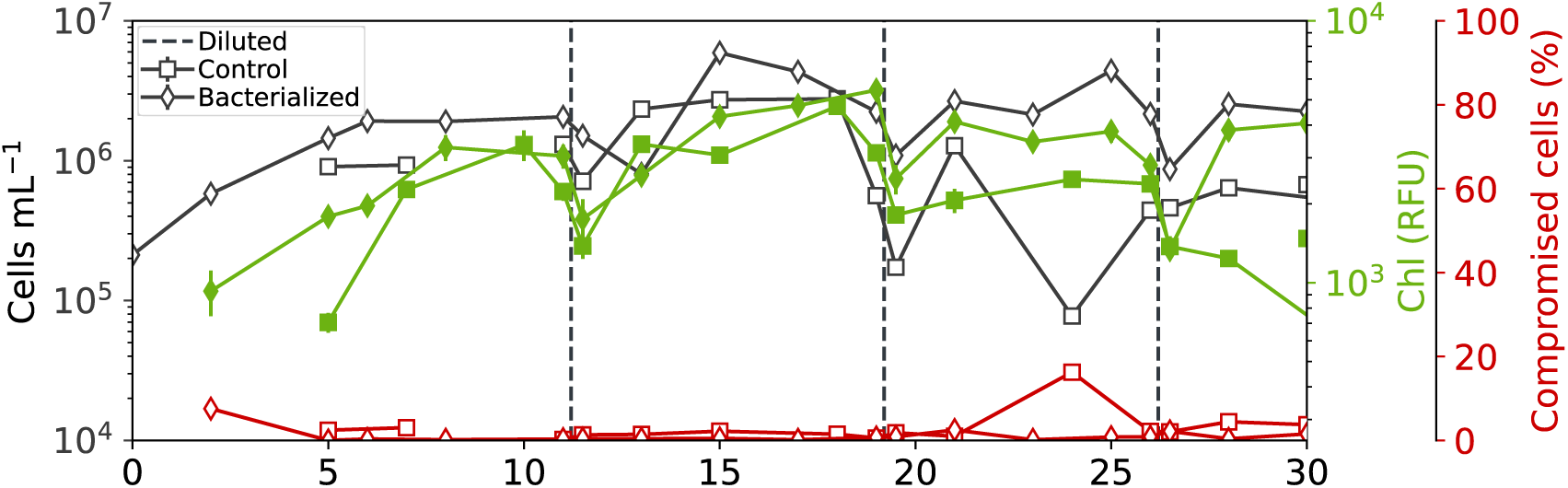
Cell count (cells mL, shown in black), chlorophyll a content (RFU, shown in green) and compromised cells (%, shown in red) for bacterialized and unbacterialized UTEX393. Dilution points are indicated with dashed lines. The unbacterialized data is indicated with squares while bacterialized data is indicated with diamonds. Each data point represents the mean value ± standard deviation (n=3).

**Figure 7:**
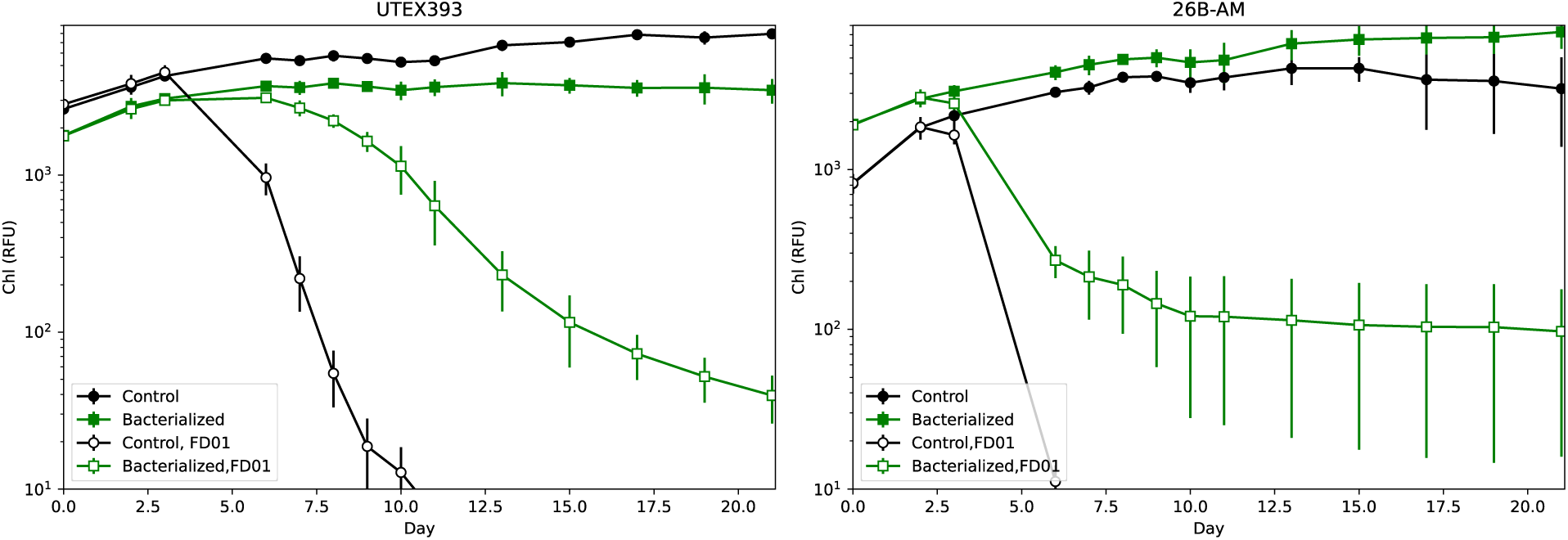
Algal biomass post-infection using chlorophyll a (RFU) as a proxy. Bacterialized cultures are shown with green squares while unbacterialized cultures are shown with black circles. Filled markers show uninfected biomass and cultures infected with amoeboaphelids are shown with open markers. Each marker indicates the mean biomass ± standard deviation (n=3).

**Figure 8:**
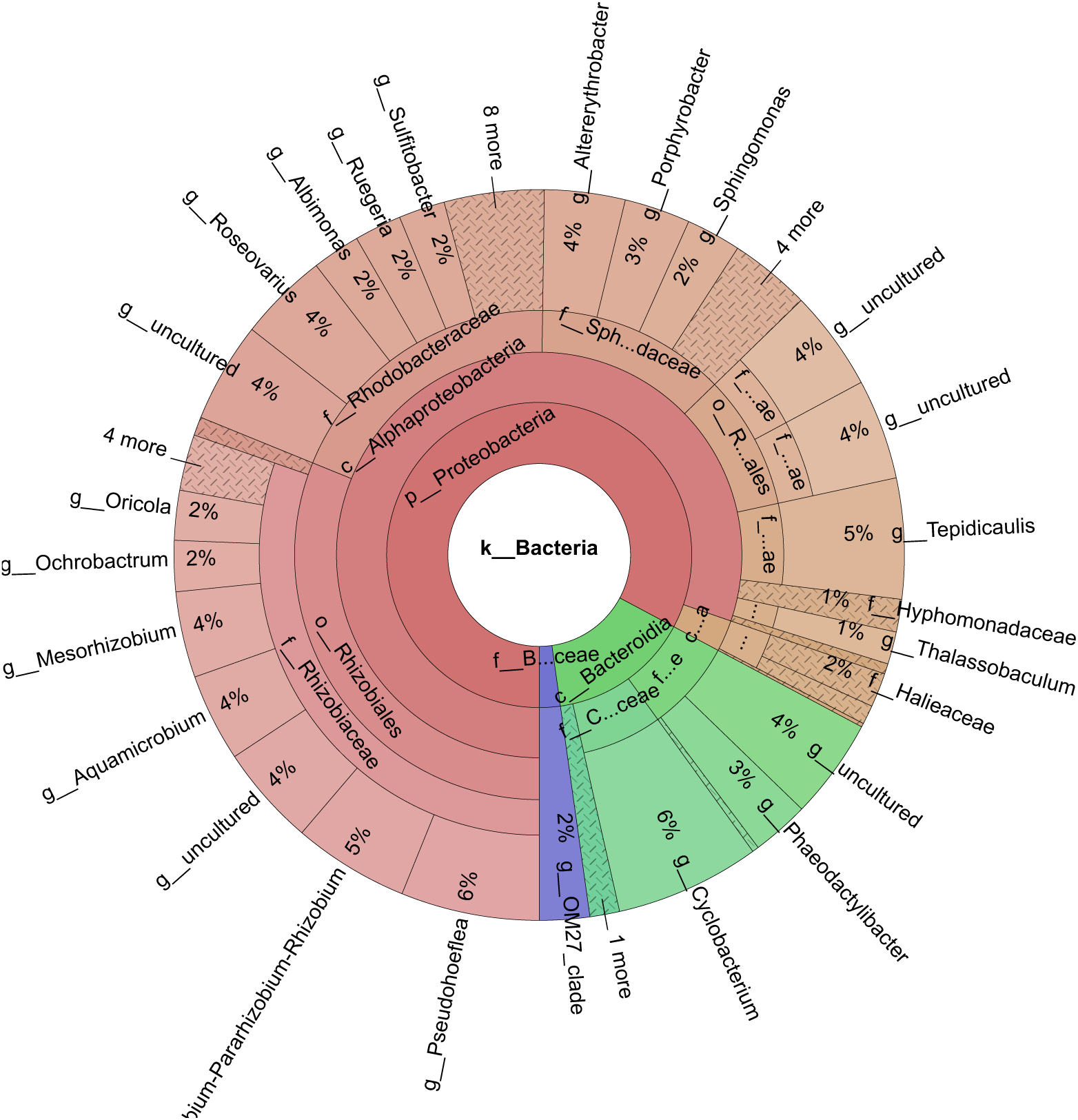
Bacterial community composition of samples taken from the most highly protected 26B-AM culture grown in an ePBR.

**Figure 9:**
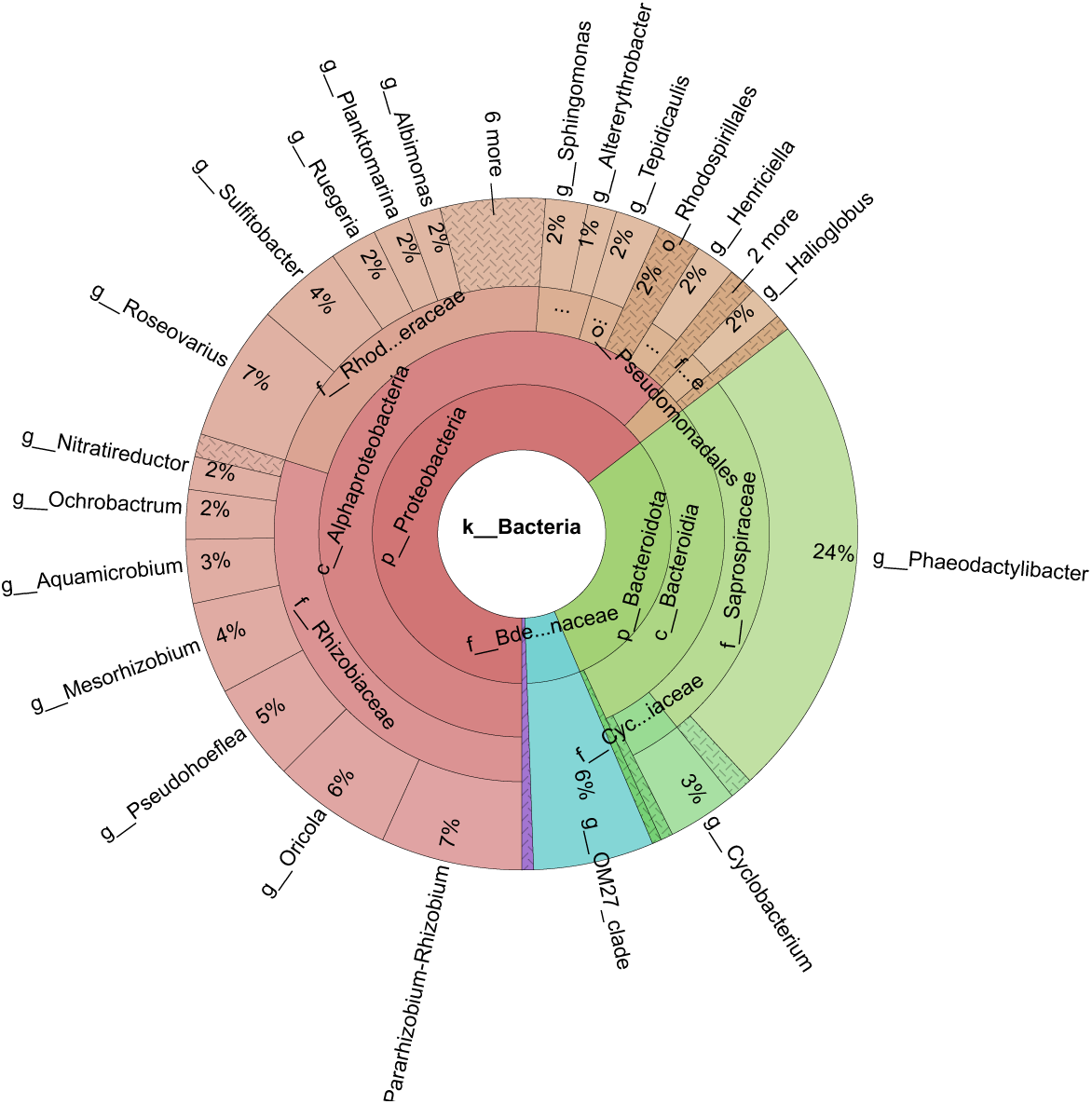
Bacterial community composition of the most protective consortium grown alongside UTEX393 in an ePBR.

Proteobacteria continue to dominate, but the largest class is now alphaproteobacteria at 80% of bacterial sequences. The largest family present is *Rhizobiales*, which contains members that live symbiotically with land plants, lichen, and algae and are responsible for nitrogen fixation in these systems [63–65]. Additionally, certain members of this family can increase algal fatty acid production [64]. Lower groups are comparatively different as well, with a more even spread in genera sequences rather than a single genus dominating the consortium.

The community composition of the most protective consortium grown with UTEX393 is less diverse than the final community grown with 26B-AM and is also less diverse than the initial bacterial consortium. It shows a somewhat lower level of protection against aphelid infection but given the lack of correlation between diversity and protection levels this is likely due to differences in community composition rather than differences in overall diversity. Like the consortium grown alongside 26B-AM, the consortium grown with UTEX393 is dominated by proteobacteria (65%), although there is a higher proportion of *Bacteroidota* in this consortium (29%). The proteobacteria are again dominated by Alphaproteobacteria (62%) and contain *Rhizobiaceae* as the dominant family (30%). Previous studies have found *Rhizobiaceae* alongside other Scenedesmus strains and other green algae species [63, 66]. *Rhizobiaceae* was not detected in the initial bacterial consortia but was detected in all algal-bacterial co-cultures, suggesting that rather than being responsible for the antifungal properties it simply grows well alongside green algae and should be expected as a member of the algal-bacterial microbiome. Rather than a single family holding responsibility for the increase in MTTF, this lack of pattern in bacterial community composition suggests again that protection against fungal infection is not due to a single bacterial genus dominating the consortium but instead may be due to the interaction between several genera as is seen in crop protection of land plants [67, 68]. Alternatively, many of the bacterial genera present are known to produce antifungal compounds [69–72] and there may be overlap between the compounds produced by different genera. While out of scope for this study, future researchers should consider not just the bacteria present but the metabolites in the system, whether algae- or bacteria-produced. However, this study demonstrated successful protection of algae by associated bacteria present in a cultivation system without the addition of external carbon sources other than the material produced by algal metabolisms and without impacting the algal productivity. This suggests that these consortia could be successfully applied in production systems without an increase in the overall biomass cost, an important step toward successful outdoor cultivation of unialgal raceway ponds for algal bioproducts. Additionally, following the identification of the mechanism of protection, microbiome engineering strategies could be used to construct a truly antifungal consortium capable of removing the aphelids from the system completely.

## 4 Conclusions

Algal biofuels have been suggested as a source of renewable energy but currently are not economically viable, partially due to their susceptibility to environmental pests. Current methods of preventing pest infection include application of costly antifungal compounds. In this study we proved the viability of instead growing algal-bacterial co-cultures that increase the MTTF of infected algal cultures by up to 350% when compared with non-bacterialized cultures. These co-cultures do not require the reapplication of bacteria but instead are stable over at least a year and show antifungal properties even as the bacterial community composition changes over time. While diversity did increase in one of the consortia over the year of culturing, there was no direct link between increased diversity and increase in MTTF. We suggest that rather than a single group holding responsibility for the antifungal properties of the consortium, there are several groups interacting to prevent infection by a common amoeboaphelid. These groups can be grown alongside the algae without the addition of bacteria-specific nutrients to the system, creating a path toward protection of outdoor unialgal raceway ponds without any additional production costs.

## 5 Supplemental Information

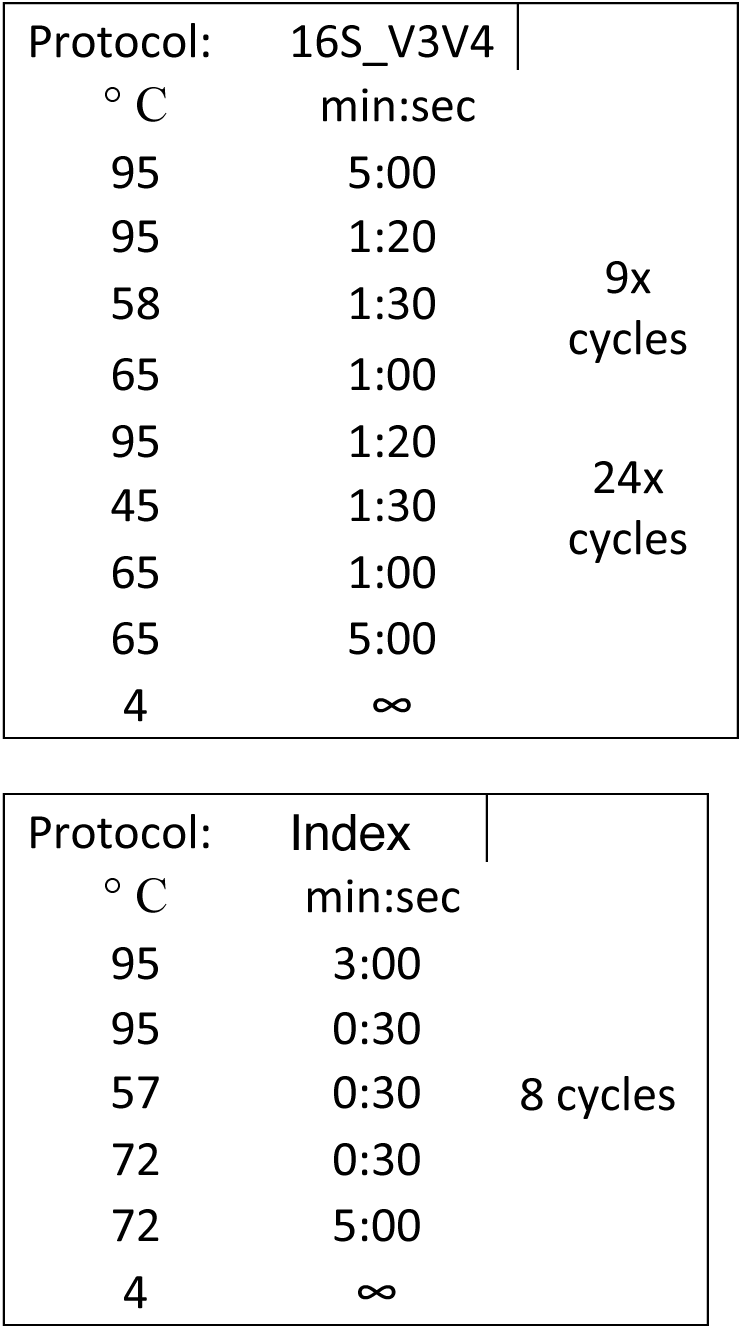

### Consortium J initial 16S sequences

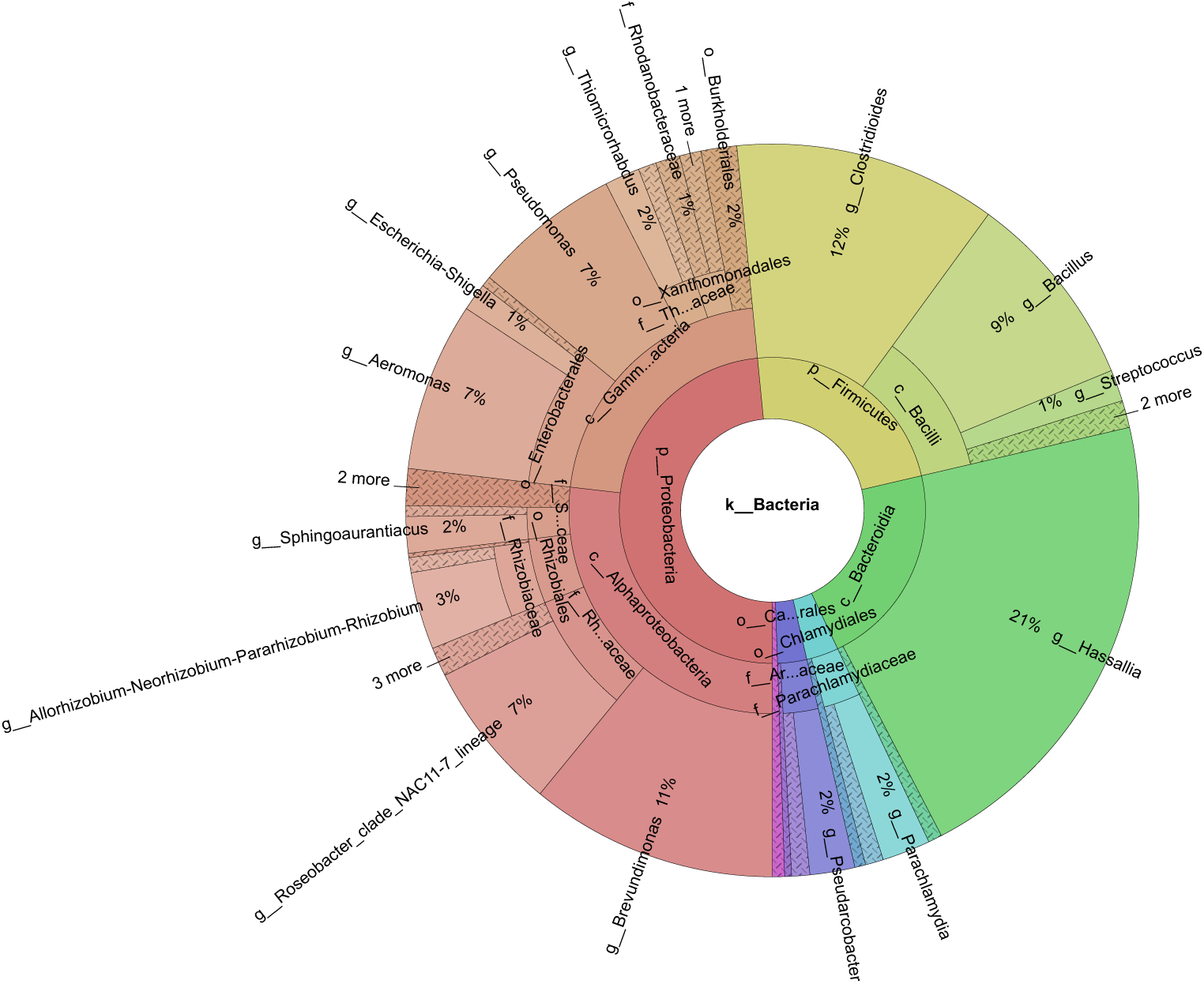

### Day 11

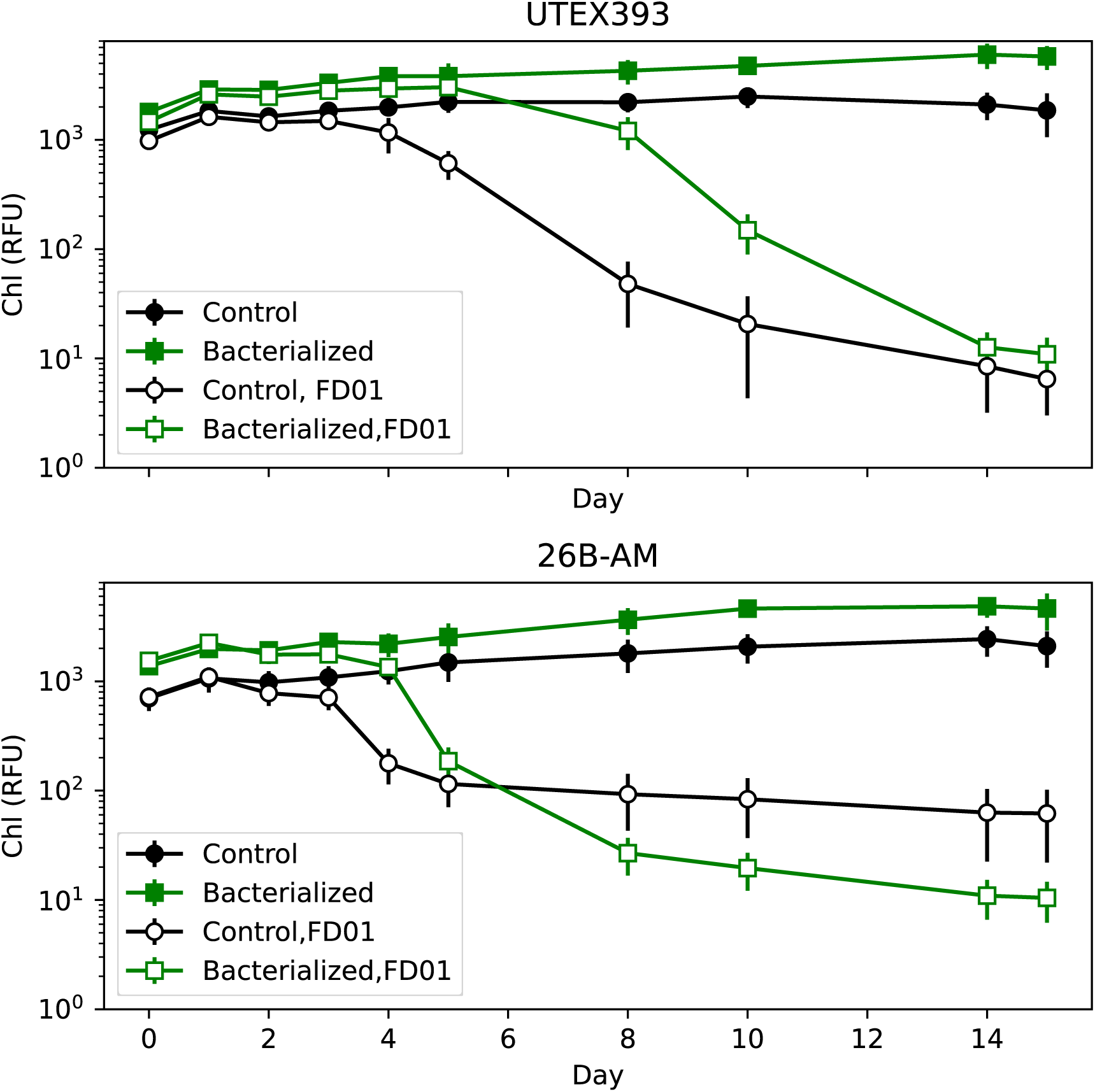

### Day 26

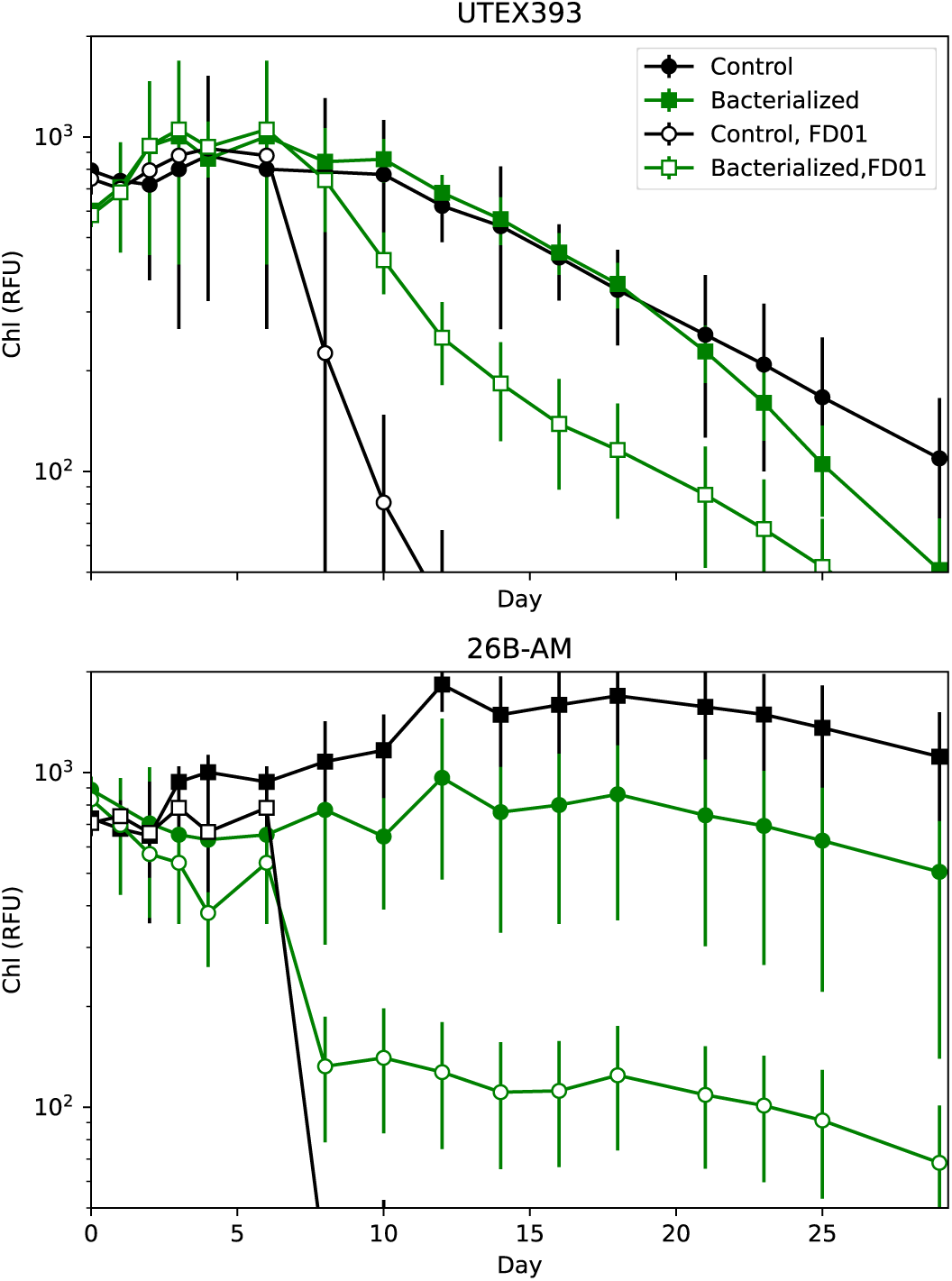

